# Opposing regulation of METTL11A by its family members METTL11B and METTL13

**DOI:** 10.1101/2022.10.05.510978

**Authors:** Haley V. Parker, Christine E. Schaner Tooley

## Abstract

N-terminal protein methylation (Nα-methylation) is a post-translational modification (PTM) that influences a variety of biological processes by regulating protein stability, protein-DNA interactions, and protein-protein interactions. Although significant progress has been made in understanding the biological roles of this PTM, we still do not completely understand how the methyltransferases that place it are regulated. A common mode of methyltransferase regulation is through complex formation with close family members, and we have previously shown that the Nα-trimethylase METTL11A (NRMT1/NTMT1) is activated through binding of its close homolog METTL11B (NRMT2/NTMT2). It has also recently been reported that METTL11A co-fractionates with a third METTL family member METTL13, which methylates both the N-terminus and lysine 55 (K55) of eukaryotic elongation factor 1 alpha (eEF1A). Here we confirm a regulatory interaction between METTL11A and METTL13 and show that, while METTL11B is an activator of METTL11A, METTL13 inhibits METTL11A activity. This is the first example of a methyltransferase being opposingly regulated by different family members. Similarly, we find that METTL11A promotes the K55 methylation activity of METTL13 but inhibits its Nα-methylation activity. We also find that catalytic activity is not needed for these regulatory effects, demonstrating new, non-catalytic functions for METTL11A and METTL13. Finally, we show METTL11A, METTL11B, and METTL13 can complex together, and when all three are present, the regulatory effects of METTL13 take precedence over those of METTL11B. These findings provide a better understanding of the regulation of Nα-methylation, and suggest a model where these methyltransferases can serve in both catalytic and non-catalytic roles.

## Introduction

METTL11A (NRMT1/NTMT1) is a distributive trimethylase belonging to the METTL (methyltransferase-like) family of proteins that methylates the N-terminus (α-amine) of its target substrates, following cleavage of the initiating methionine (1,2). METTL11A methylates both canonical and non-canonical consensus sequences. The canonical X-P-K sequence allows Ala, Pro, Ser, Gly, or Met in the first position and requires Pro and Lys in the second and third, respectively (1). The non-canonical sequence expands to include one of seven amino acids other than Pro in the second position (A/S/G/M/E/N/Q), and either Lys or Arg in the third (3). Combined, the two consensus sequences predict over 300 METTL11A targets, and verified substrates include regulator of chromatin condensation 1 (RCC1), zinc fingers and homeoboxes 2 (ZHX2), and the ribosomal proteins RPS25 and RPL12 (1,4,5).

Our lab and others have identified many downstream effects of the loss of METTL11A including impaired DNA damage repair, altered cell proliferation, abnormal muscle cell differentiation, and premature neural stem cell depletion (6-9). METTL11A acts as a tumor suppressor in breast cancer cells, as its knockdown promotes DNA damage, cell proliferation, invasive potential, and xenograft tumor growth (6). METTL11A acts as an oncogene in colon and cervical cancer, as its loss slows growth and reduces invasion and migration capability, respectively (10,11). CRISPR-Cas9 mediated METTL11A knockout in C2C12 myoblasts prevents their differentiation into myofibers and instead promotes osteoblastic phenotypes (8). Knockout of *Mettl11A* in mice results in premature ageing phenotypes, including depletion of the neural stem cell pools, neurodegeneration, and cognitive impairments (9). Despite identifying these important biological roles of METTL11A, we still do not have a thorough understanding of its upstream regulation.

One form of METTL11A regulation that we have identified is through physical interaction with its homolog, METTL11B (NRMT2/NTMT2) (12). Using analytical ultracentrifugation, METTL11A was found to primarily exist as a dimer, and when combined with METTL11B, formed a heterotrimer consisting of the METTL11A dimer bound to a METTL11B monomer (12). Through this interaction, METTL11B stabilizes METTL11A and specifically promotes METTL11A methylation of non-canonical targets (12). As METTL11B primarily acts as a Nα-monomethylase *in vitro*, we hypothesized METTL11B could be performing the first methylation event and priming METTL11A substrates for subsequent di- and trimethylation (2). However, a catalytically dead mutant of METTL11B was also capable of activating METTL11A (12), suggesting that METTL11A activation is a non-catalytic function of METTL11B.

Similar regulatory interactions are apparent among other methyltransferase pairs, including the RNA methyltransferases METTL3 and METTL14, as well as the DNA methyltransferases DNMT3A, DNMT3B, and DNMT3L (13-21), where often one member of these pairs is serving a non-catalytic function. METTL3 is the catalytic component of the complex responsible for writing N^6^-methyladenosine (m^6^A), while the primary role of METTL14 is to provide structural support and aid in substrate recognition to promote METTL3 methylation activity (13-16). METTL14 possesses a relatively occluded active site, suggesting it does not carry out catalytic activity of its own, but instead has an important non-catalytic function as an allosteric regulator of METTL3 (14-16). Similarly, compared to DNMT3A and DNMT3B, DNMT3L lacks important motifs in the catalytic site, rendering it catalytically inactive, but still able to provide structural support to DNMT3A and DNMT3B to stimulate their methylation activities (17-26).

Interestingly, METTL11A was reported to co-fractionate with another member of the METTL family, METTL13 (27). METTL13 is a dual-function methyltransferase that methylates eukaryotic elongation factor 1 alpha (eEF1A) on its Nα-amine and on internal lysine 55 (K55) (28,29). eEF1A, which is the collective reference to paralogous proteins eEF1A1 and eEF1A2, is a GTP-binding protein that is vital for the elongation step of translation (30-32). METTL13 contains two distinct 7β strand methyltransferase domains, with the N-terminal methyltransferase domain (NTD) targeting K55 of eEF1A, and the C-terminal methyltransferase domain (CTD) targeting its N-terminus (28,29). Aberrant expression of METTL13 is linked to cancer, although like METTL11A, the mechanism of oncogenesis appears to be cell type specific. In clear cell renal cell carcinoma and bladder cancer, METTL13 has been found to inhibit cancerous phenotypes and is associated with more favorable patient outcomes (33,34). However, upregulation of METTL13 and K55me2 eEF1A is associated with poor prognosis in lung and pancreatic cancer patients (29).

Here we confirm the interaction between METTL11A and METTL13, and demonstrate for the first time the regulatory nature of this interaction. In contrast to METTL11B, METTL13 inhibits METTL11A methylation of both canonical and non-canonical substrates. Reciprocally, we show that METTL11A can both inhibit METTL13 methylation of the N-terminus and promote methylation of K55, and catalytic activity is not needed for the regulatory roles of either METTL11A or METTL13. Finally, we demonstrate that METTL11A, METTL11B, and METTL13 can complex together, and when this occurs, regulation of METTL11A by METTL13 takes precedence over that of METTL11B. Our results not only describe the first example of a methyltransferase participating in opposing regulatory interactions with close family members, but also identify novel non-catalytic, regulatory functions of both METTL11A and METTL13. These findings on the regulatory mechanisms of Nα-methylation will help us to better understand how dysregulation can lead to the development of disease.

## Results

### Identification of residues that regulate the METTL11A/METTL11B interaction

We have previously identified that METTL11A and METTL11B interact to form a heterotrimer composed of a METTL11A dimer and a METTL11B monomer (12). Through this interaction, METTL11B provides stability to METTL11A and enhances METTL11A trimethylation activity on non-canonical substrates (12). Using computational modeling, we previously published a model of the METTL11A/METTL11B interaction and proposed twelve pairs of residues from METTL11A and METTL11B that are predicted to be important for mediating this interaction (12). Here, we first aimed to verify this model by mutating predicted residues and assaying the effect on METTL11A binding to METTL11B.

From the list of twelve predicted pairs, we prioritized mutations that also have biological relevance. Using the Catalogue of Somatic Mutations in Cancer (COSMIC), we found three METTL11B cancer-associated mutations (Q67H-lung, F68L-colon, D232N-prostate) that affected residues predicted to be important for binding from our previous model (12,35). These three mutations, and a fourth cancer-associated METTL11B mutation (V224L-breast) that is catalytically inactive but does not mediate binding to METTL11A, were all introduced individually into human METTL11B-GFP. Either WT or mutant METTL11B-GFP was transiently transfected into human embryonic kidney (HEK293T) cells along with human METTL11A-FLAG, and co-immunoprecipitations (co-IPs) were performed. Western blots were used to compare the ability of WT and mutant METTL11B-GFP to co-IP with METTL11A-FLAG (Fig. 1A). Only METTL11B-GFP possessing the cancer-associated D232N mutation showed a significant decrease in its ability to co-IP with METTL11A-FLAG as compared to WT METTL11B-GFP (Fig. 1, A and B). No other mutations had a significant effect on the METTL11A/METTL11B interaction (Fig. 1B). These results suggest that D232 of METTL11B is an important residue for the interaction between METTL11A and METTL11B.

**Figure 1.**
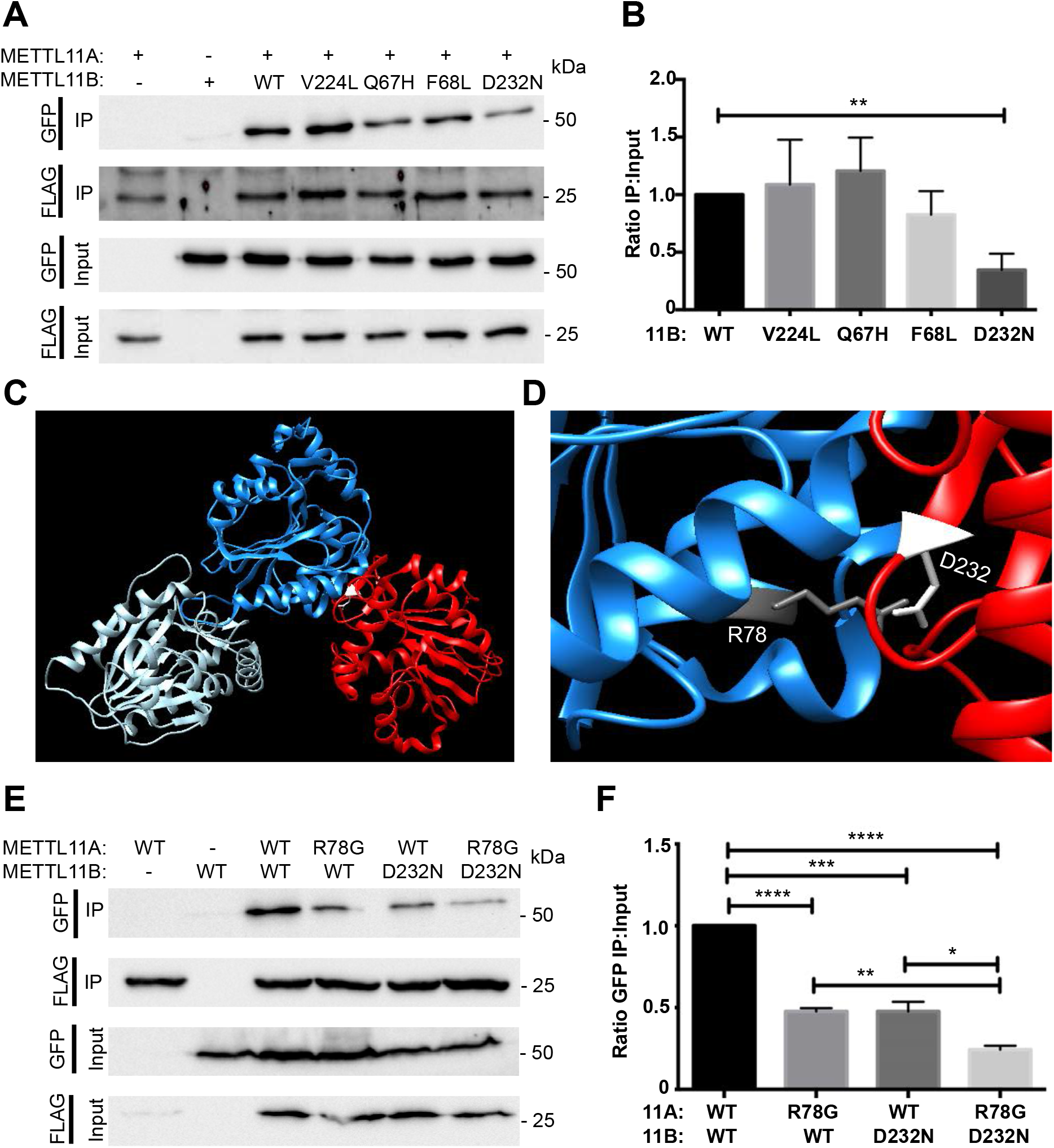
Predicted mutations regulate the METTL11A/METTL11B interaction. *A*, An Asp residue in METTL11B (D232), predicted by modeling to regulate binding to METTL11A (12) and found mutated to Asn in prostate cancer (D232N) (35), disrupts the ability of METTL11B-GFP to co-IP with METTL11A-FLAG. *B*, Quantification of blots showing the ratio of pulled-down METTL11B-GFP to GFP input. *C*, ZDOCK model of the METTL11A/METTL11B interaction with METTL11A dimer shown in light and dark blue and METTL11B monomer shown in red. *D*, The model predicts METTL11A R78 as the interacting partner of METTL11B D232. *E*, Expression of both METTL11B D232N and METTL11A R78G further disrupts binding. *F*, Quantification of blots showing the ratio of pulled-down METTL11B-GFP to GFP input. These data indicate disease mutations can disrupt METTL11A/METTL11B interactions. n=3, * denotes p<0.05, ** denotes p<0.01, *** denotes p<0.001, **** denotes p<0.0001 as determined by unpaired t-test.

Having positively identified the importance of METTL11B D232, we next aimed to update our computational model with D232 included as a required element of the interaction surface. Docking-based modeling was used to predict the new protein-protein interaction interface that results from this added constraint. Computational models were generated with the ZDOCK server and graphics were viewed and analyzed using Chimera (36,37). This method was first used to generate a model of the METTL11A (PDB 5E1B) dimer that closely correlated with our previously published METTL11A dimer model (12). The METTL11A dimer model was then docked to a METTL11B (PDB 6DUB) monomer using the ZDOCK server, with METTL11B D232 identified as a contacting residue (Fig. 1C). The resulting new model of the METTL11A/METTL11B interaction shared many similarities with our previously published model (12). The updated model suggested METTL11B D232 interacts with METTL11A R78 (Fig. 1D). To verify if METTL11A R78 is also important for the METTL11A/METTL11B interaction, we next repeated the METTL11A-FLAG/METTL11B-GFP co-IPs with METTL11A R78G alone or in combination with METTL11B D232N. Similar to METTL11B D232N alone, METTL11A R78G alone also significantly reduced the interaction between METTL11A and METTL11B (Fig. 1, E and F). When both mutants were expressed together, there was an even greater decrease in the interaction (Fig. 1, E and F), confirming the importance of this residue pair, showing that both members play important regulatory roles, and further refining our interaction model.

### METTL11A also interacts with METTL13

Since other interacting methyltransferases, including METTL3/METTL14, also exist as part of larger complexes (reviewed in (38-42)), we were interested in identifying additional METTL11A/METTL11B binding partners. A study completed by Havugimana et al. previously identified that METTL11A co-fractionated with another METTL family member, METTL13 (27). To verify METTL11A and METTL13 interact, we again performed co-IP experiments. HEK293T cells were transfected with METTL11A-FLAG and METTL13-GFP, and western blots determined that METTL13-GFP did co-IP with METTL11A-FLAG (Fig. 2A). This was interesting to us, as we have previously shown by immunofluorescence that METTL11A is predominantly nuclear and inactive towards RCC1 in the cytoplasm (1,3), and METTL13 is thought to be primarily cytoplasmic (43). To better understand the cell compartment localization of METTL11A and METTL13, we performed nuclear/cytoplasmic fractionations and found that while METTL13 is predominantly cytoplasmic, METTL11A is found in both the nucleus and cytoplasm (Fig. 2B). These data indicate METTL11A and METTL13 are interacting in the cytoplasm. To confirm these findings, HEK293T cells were transfected with METTL13-FLAG, cell lysates were fractionated into cytoplasmic and nuclear fractions, METTL13-FLAG was immunoprecipitated out of the cytoplasmic fraction, and interactors were identified using liquid chromatography-mass spectrometry (LC-MS). Six proteins were identified as having a 100% abundance ratio in the METTL13-FLAG transfected cells as compared to the untransfected control cells (Fig. 2C). Of these six proteins, METTL11A had the highest percent coverage and number of representative peptides (Fig. 2C), further verifying the METTL11A/METTL13 interaction.

**Figure 2.**
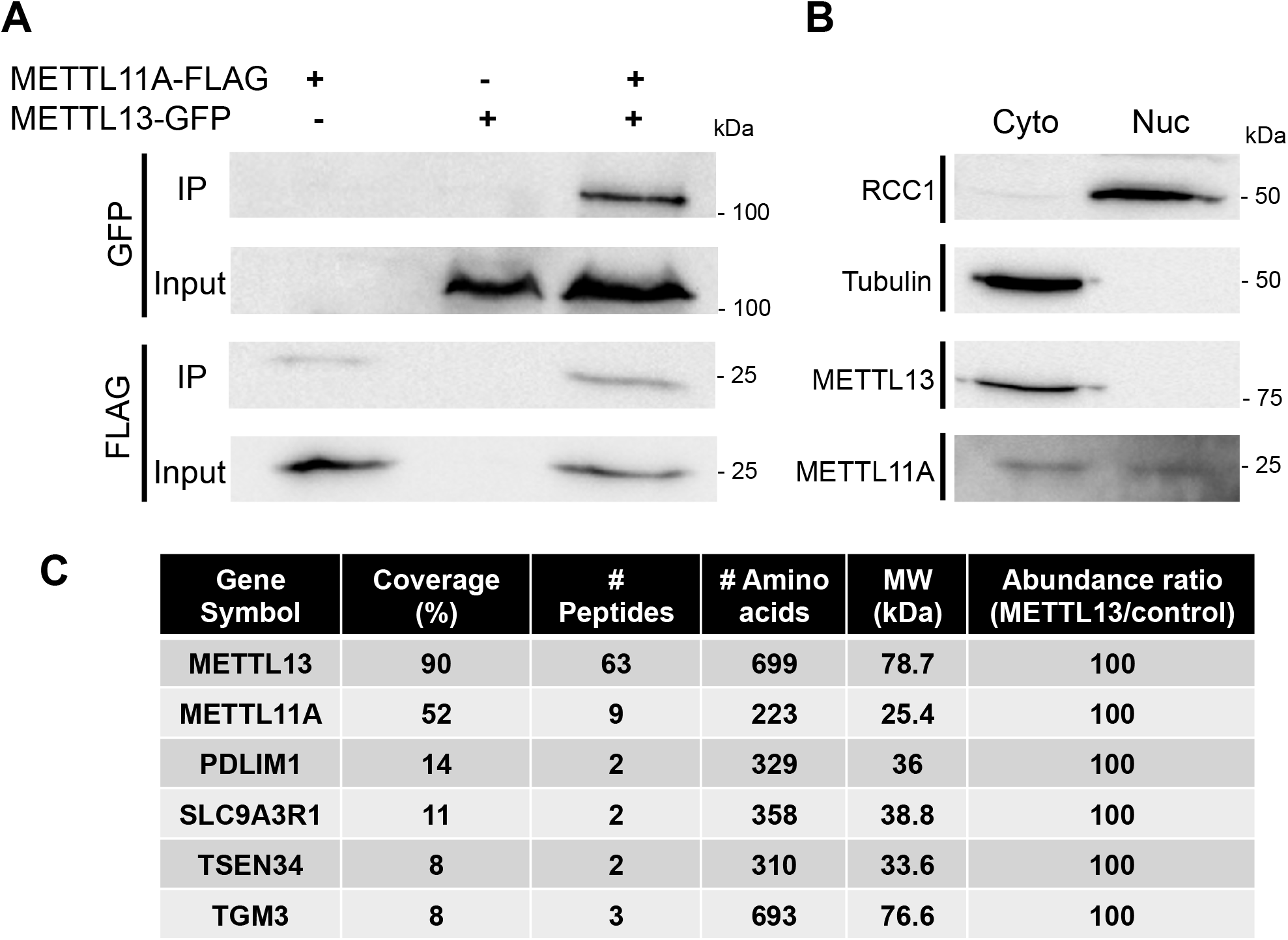
METTL13 and METTL11A interact. *A*, METTL13-GFP co-IP’s with METTL11A-FLAG. *B*, Untransfected HEK293T cellular fractionation show endogenous METTL13 primarily in the cytoplasmic fraction and endogenous METTL11A in both the nuclear and the cytoplasmic fractions. *C*, Top six proteins pulled down with METTL13-FLAG in the cytoplasm. These data verify a cytoplasmic interaction between METTL11A and METTL13.

### METTL11A and METTL13 exhibit reciprocal regulation

As the interaction between METTL11A and METTL11B results in the activation of METTL11A methylation activity, we next determined if METTL13 could similarly regulate METTL11A activity. *In vitro* methyltransferase assays were performed using combinations of recombinant human METTL11A and METTL13 enzymes and an RCC1 N-terminal peptide as substrate. We found that METTL13 inhibited METTL11A trimethylation activity of RCC1 (Fig. 3, A and B). Specifically, METTL13 was able to both significantly increase the Km of METTL11A and significantly lower its Vmax (Fig. 3, C and D). These data indicate METTL13 is a mixed inhibitor of METTL11A and it regulates METTL11A in an opposing manner to METTL11B.

**Figure 3.**
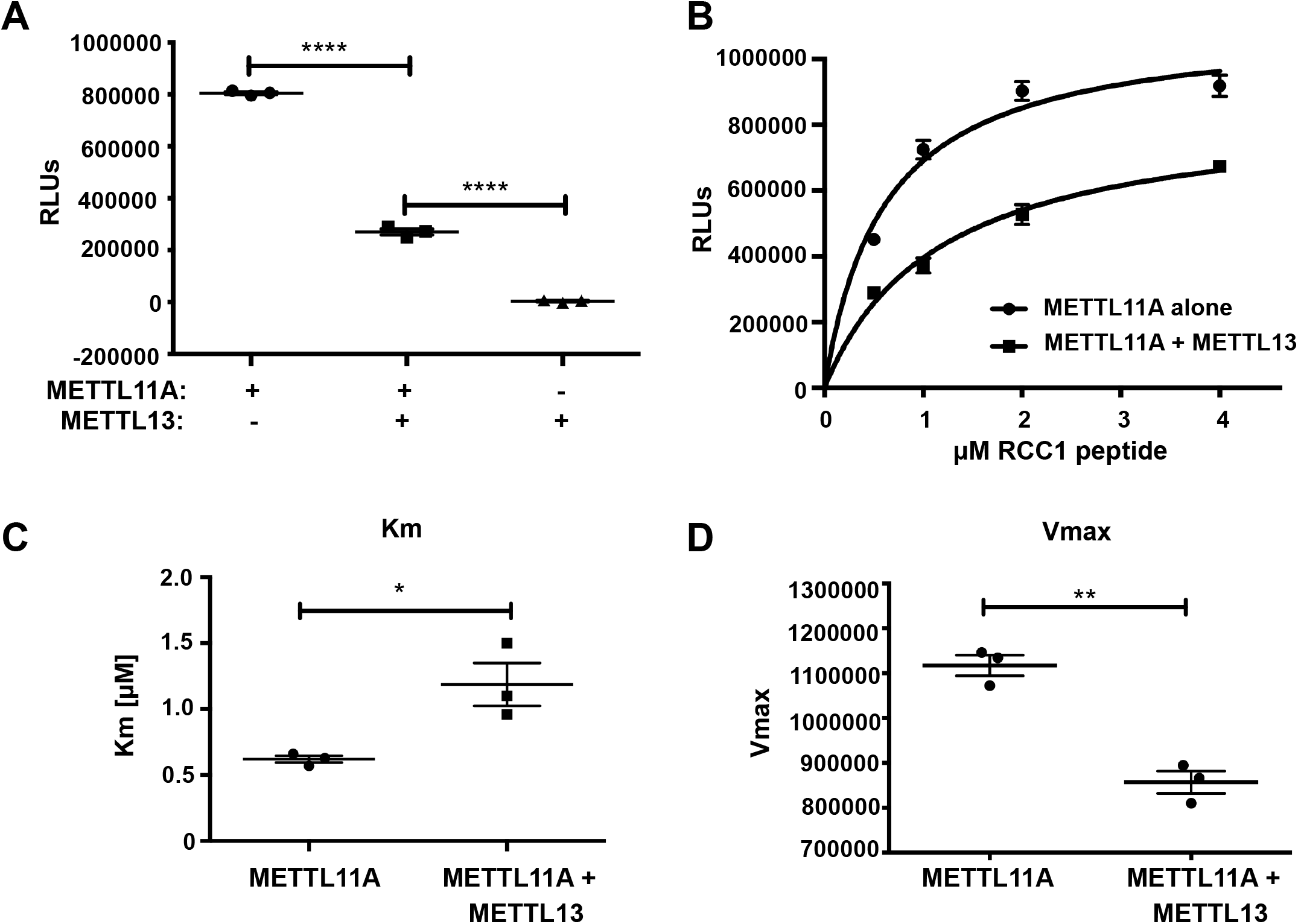
METTL13 inhibits the ability of METTL11A to methylate RCC1. *A*, Methylation of an RCC1 N-terminal peptide is significantly lower when METTL13 is present compared to METTL11A by itself. *B*, Activity curves of RCC1 methylation by METTL11A in the presence or absence of METTL13. *C*, Km is significantly higher when METTL13 is present. *D*, Vmax is significantly lower when METTL13 is present. These data indicate METTL13 is a mixed inhibitor of METTL11A. n=3, * denotes p<0.05, ** denotes p<0.01, **** denotes p<0.0001 as determined by unpaired t-test.

We were next interested in examining if METTL11A can alter METTL13 activity. METTL13 is a dual-function methyltransferase that can methylate both the N-terminus and internal lysine 55 (K55) of eEF1A. We first used *in vitro* methyltransferase assays with an N-terminal eEF1A peptide substrate to measure the effect of METTL11A on the ability of METTL13 to Nα-methylate eEF1A. We found that METTL11A exhibited a partial, though significant, inhibition of METTL13 Nα-methylation of eEF1A at high concentrations (Fig. 4, A and B). Unlike the effect of METTL13 on METTL11A, we found a significant decrease in Vmax but no significant difference in Km (Fig. 4, C and D), indicating METTL11A is acting as a non-competitive inhibitor of METTL13 Nα-methylation of eEF1A.

**Figure 4.**
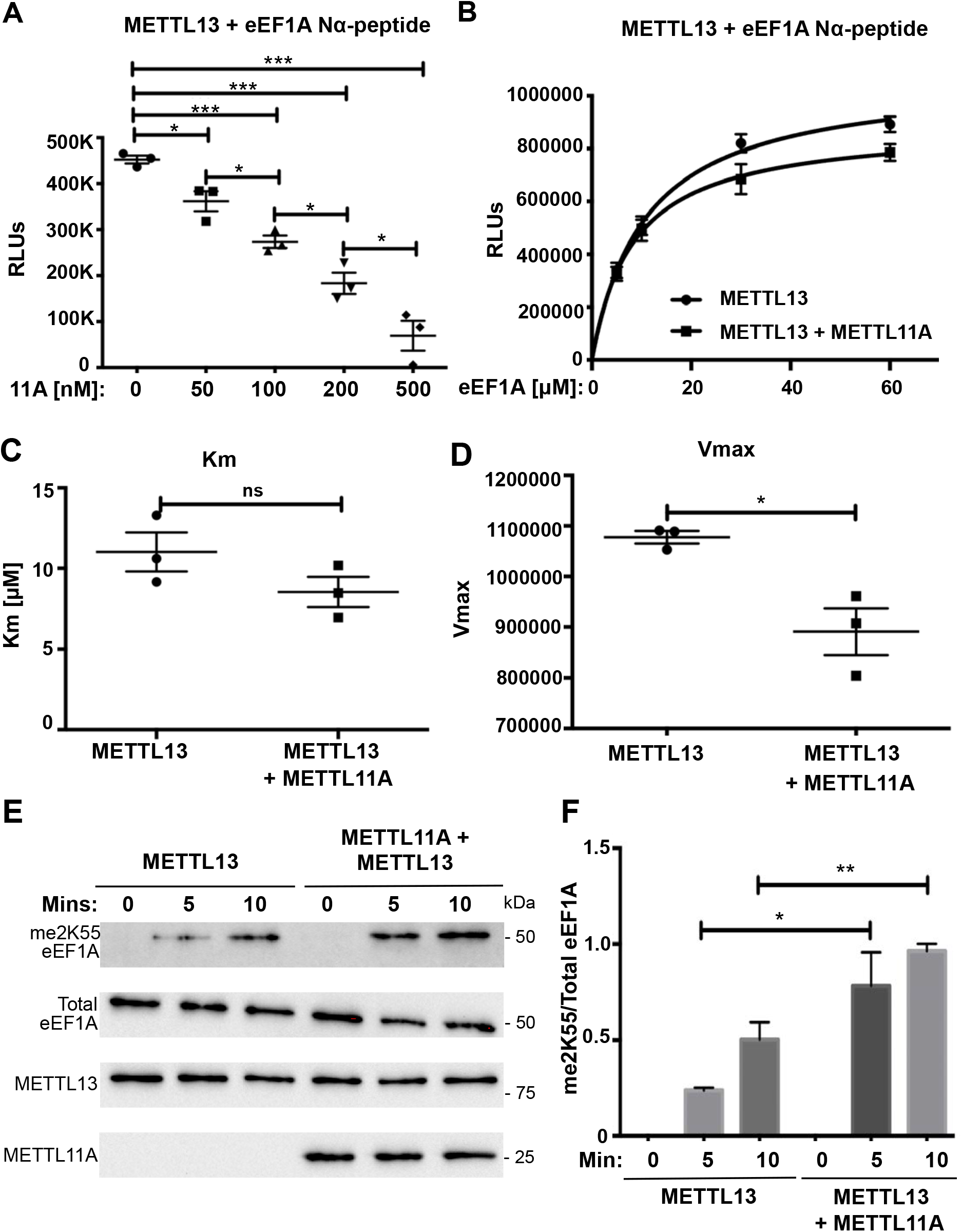
METTL11A inhibits the ability of METTL13 to methylate eEF1A at the N-terminus but promotes methylation at K55. *A*, Methylation of an eEF1A N-terminal peptide is significantly lower as METTL11A concentration increases compared to METTL13 by itself. *B*, Activity curves of eEF1A Nα-methylation by METTL13 in the presence or absence of METTL11A. *C*, Km is not significantly different when METTL11A is present. *D*, Vmax is significantly lower when METTL11A is present. *E*, Levels of dimethylated lysine 55 (me2K55 eEF1A) are higher at 5 and 10 minutes when METTL11A is present with METTL13 compared to METTL13 alone. *F*, Quantification of blots showing the ratio of me2K55 eEF1A to total eEF1A. These data indicate METTL11A is a non-competitive inhibitor of eEF1A Nα-methylation and an activator of K55 methylation. n=3, * denotes p<0.05, ** denotes p<0.01, *** denotes p<0.001 as determined by unpaired t-test.

We then tested if METTL11A affects the ability of METTL13 to methylate K55. Unfortunately, it has been shown that peptides containing K55 are not viable substrates for *in vitro* methyltransferase assays (29). It has been suggested that the methyltransferase domain responsible for methylating K55 may require the substrate to be in its fully folded state rather than an unstructured short peptide (29). Thus, we used full-length, recombinant eEF1A protein substrate and K55 methyl-specific antibodies to assay METTL13 activity by western blot (Fig. 4E). We found a significant increase in eEF1A K55 dimethylation by METTL13 at both 5 and 10 min when METTL11A is present compared to METTL13 alone (Fig. 4F). Together, these results suggest that while METTL13 decreases the methylation activity of METTL11A, METTL11A causes METTL13 to favor eEF1A K55 methylation over Nα-methylation. Further functional studies will be needed to further delineate the implications of this shift, as the distinct roles of each methylation remain unknown.

### Non-catalytic activities of METTL11A and METTL13

The regulation of METTL11A is a non-catalytic function of METTL11B, and another family member, METTL16, has also recently been shown to have a non-catalytic role facilitating assembly of the translation-initiation complex, in addition to its role as an m^6^A methyltransferase (44). To determine if the regulatory roles of METTL11A and METTL13 are dependent on their catalytic activities, we repeated the previous *in vitro* methyltransferase assays with catalytically inactive mutant METTL13 and METTL11A. The METTL13 mutations G58R and E524A were previously found to abrogate methylation activity of the METTL13 NTD and CTD, respectively (28,29). We constructed a G58R/E524A double mutant and tested its ability to inhibit METTL11A methylation of RCC1. Although lacking significant catalytic activity of its own (Fig. S1, A and B), the G58R/E524A METTL13 double mutant was able to inhibit METTL11A methylation of RCC1 at a level similar to that of WT METTL13 (Fig. 5A). Mutation of Asp180 to Lys or Asn (D180K/D180N) in METTL11A has also been shown to inhibit its catalytic activity (1,45). Similar to the METTL13 double mutant, D180K METTL11A was able to inhibit METTL13 Nα-methylation of eEF1A at levels comparable to WT METTL11A (Fig. 5B), despite lacking catalytic activity of its own (Fig. S1C). Together these results demonstrate novel, non-catalytic regulatory functions for both METTL11A and METTL13.

**Figure 5.**
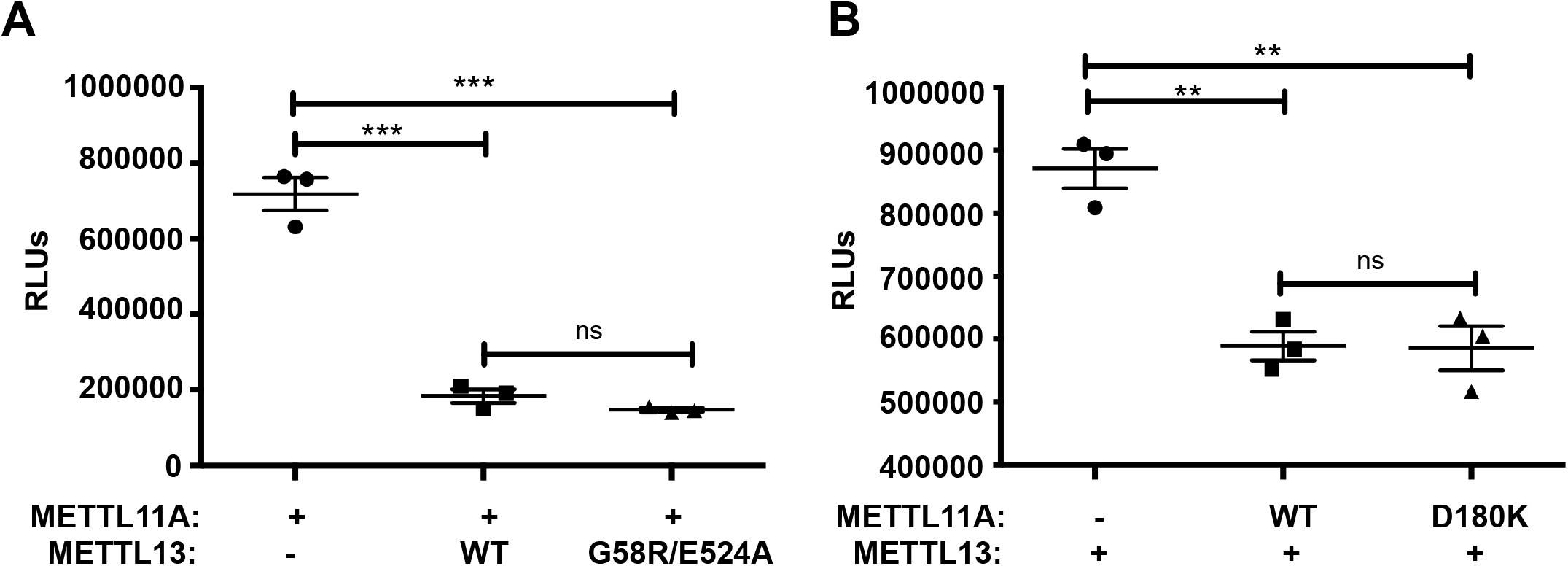
Regulatory effects of METTL11A and METTL13 are not dependent on their catalytic activities. *A*, Catalytically inactive METTL13 possessing the G58R and E524A mutations inhibits METTL11A methylation of the RCC1 peptide at a comparable level to WT METTL13. *B*, Catalytically inactive METTL11A possessing the D180K mutation inhibits METTL13 methylation of the eEF1A N-terminal peptide at a comparable level to WT METTL11A. These data indicate the catalytically inactive METTL13 and METTL11A can still elicit their regulatory effects. n=3, ** denotes p<0.01, *** denotes p<0.001 as determined by unpaired t-test.

### METTL11A/METTL11B/METTL13 interactions

Once we determined that the METTL11A/METTL13 interaction exhibits regulatory effects on both members of the pair and the regulatory effects were likely structural (not catalytic), we wanted to perform computational modeling of the interaction interface. However, METTL13 is relatively large (∼79 kDa) and contains two distinct methyltransferase domains, each responsible for targeting one of the methylation sites on eEF1A (28). To determine which region of METTL13 was responsible for mediating the interaction with METTL11A, we created GFP-tagged N-terminal and C-terminal fragments of METTL13 (Fig. 6A). The N-terminal fragment (residues M1-Y344) contained the methyltransferase domain that targets K55 of eEF1A, and the C-terminal fragment (residues E345-V699) contained the methyltransferase domain that targets the N-terminus of eEF1A (28). Each was expressed in HEK293T cells with METTL11A-FLAG and co-IPs were performed. Only the N-terminal METTL13 fragment was found to co-IP with METTL11A-FLAG (Fig. 6A), suggesting that residues M1-Y344 interact with METTL11A. This suggests that binding at one region of METTL13 can affect activity at both domains. The currently available crystal structure of METTL13 only consists of the C-terminal domain (residues C470-V699) (28), so we first used the I-TASSER server to predict a model of METTL13 residues M1-Y344 (46-48). This METTL13 N-terminal fragment model was then used as input for the ZDOCK server (36), with the previously described METTL11A dimer model, to obtain a model of the predicted METTL13/METTL11A interaction interface (Fig. 6B). Nine amino acid pairs were predicted to regulate this interaction (Fig. 6C).

**Figure 6.**
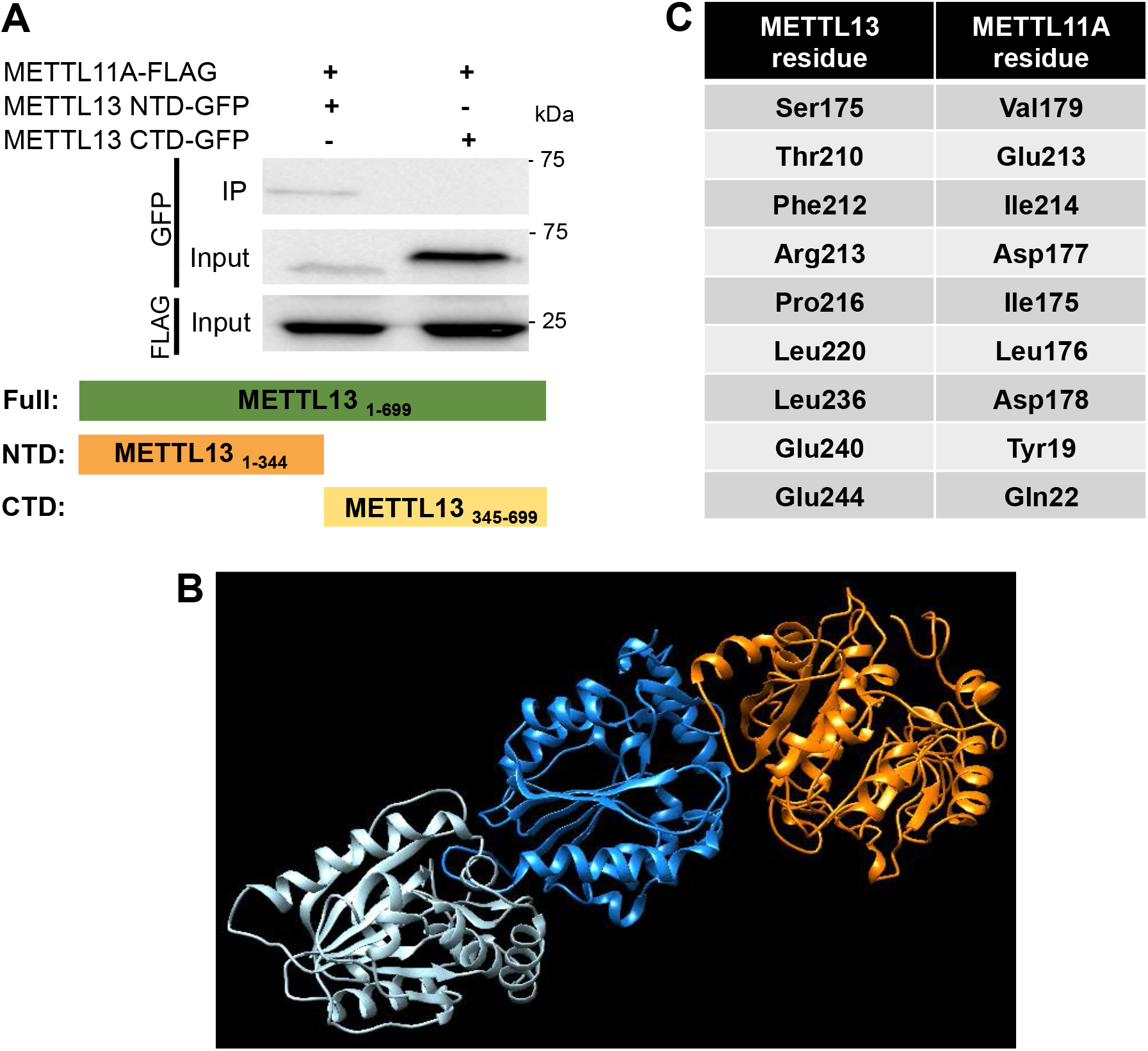
METTL13 interacts with METTL11A through its N-terminal domain. *A*, METTL11A interacts with the N-terminal domain (NTD) of METTL13 (residues 1-344), which has eEF1A K55 methylation activity. *B*, ZDOCK model of the METTL11A dimer (shown in shades of blue) bound to the METTL13 NTD (model obtained using the I-TASSER server; shown in orange). *C*, This model predicts that nine pairs of amino acids mediate the METTL11A/METTL13 interaction.

Since we had now determined that METTL11A is involved in opposing regulatory interactions with two different METTL family members, METTL11B and METTL13, we were next interested in determining if METTL11A, METTL11B, and METTL13 can exist in a complex together or if the interactions are mutually exclusive. We first used co-IPs to determine that METTL13 and METTL11B can interact (Fig. 7A), indicating they could be in a complex together. To determine if this interaction was dependent on METTL11A acting as a bridge between METTL11B and METTL13, we performed similar experiments in control HCT116 cells and HCT116 cells with CRISPR/Cas9-mediated knockout of METTL11A (10). We found that the interaction between METTL13 and METTL11B is enhanced when METTL11A is present in control cells compared to in the METTL11A KO cells (Fig. 7, A and B), indicating they can all be in a complex together. When the previously described predicted interaction interfaces for METTL11A with either METTL11B (Fig. 1C) or METTL13 (Fig. 6B) are shown simultaneously on a space-filling model of the METTL11A dimer, one of the METTL13 binding interfaces overlaps with the METTL11A dimer interface, indicating only one METTL13 monomer can bind the METTL11A dimer (Fig. 7C). The remaining METTL13 interface is in close proximity to one of the METTL11B interfaces, suggesting that the binding of METTL13 would preclude the binding of METTL11B at that site, leaving the single remaining METTL11B interface available for binding (Fig. 7C). This model is consistent with the previously determined 2:1 ratio of METTL11A:METTL11B binding (12).

**Figure 7.**
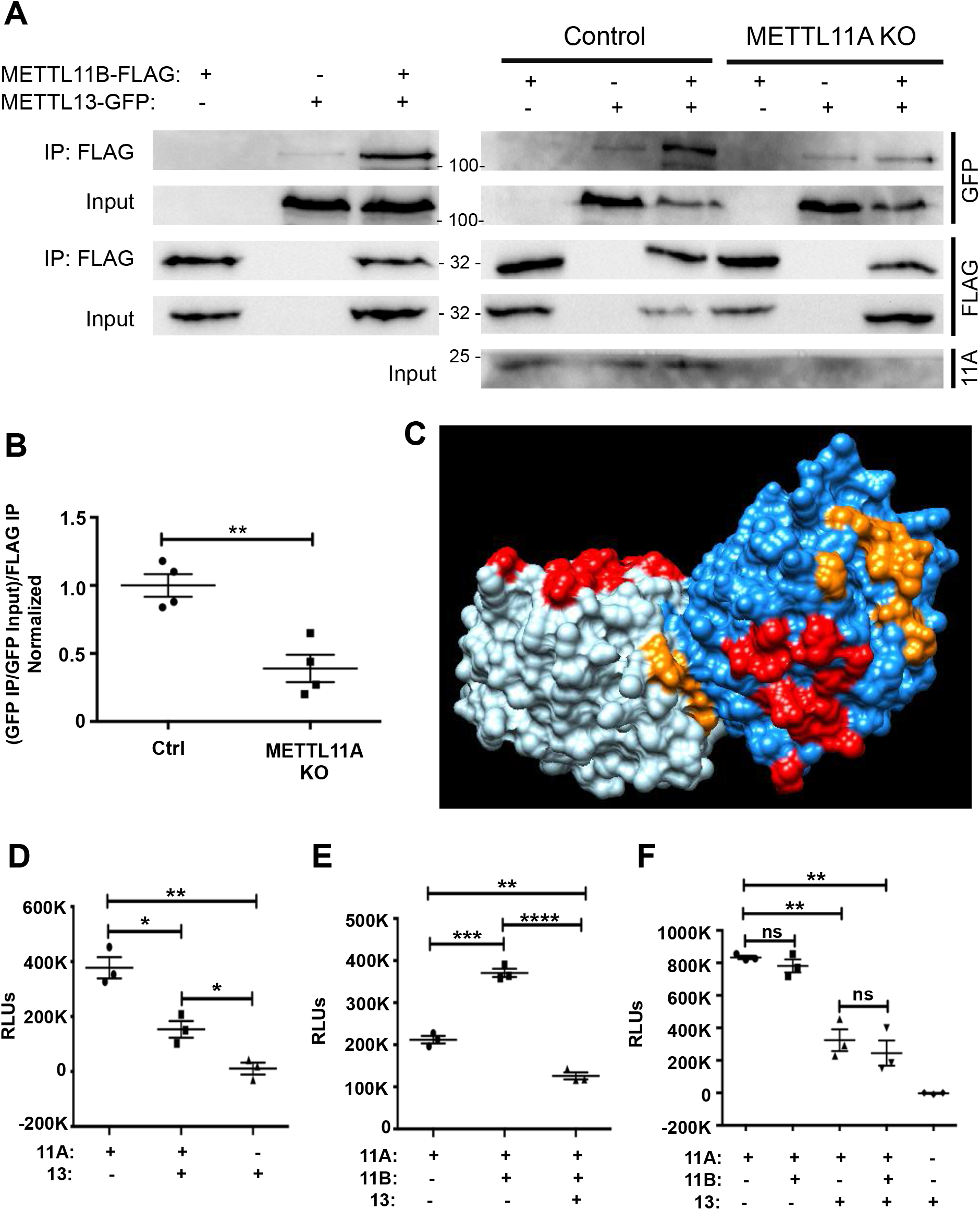
METTL13 outcompetes METTL11B for regulation of METTL11A. *A*, Left - In HEK293T cells, METTL13-GFP co-IPs with METTL11B-FLAG. Right - METTL13-GFP co-IPs with METTL11B-FLAG better when METTL11A is present in control HCT116 cells compared to METTL11A KO HCT116 cells. *B*, Quantification of blots showing the ratio of pulled-down METTL13-GFP to GFP input, ratioed to pulled-down METTL11B-FLAG. Values were normalized to 1. n=4. *C*, Model of METTL11A dimer (shown in light and dark blue), with predicted METTL13 interaction interfaces shown in orange and predicted METTL11B interaction interfaces shown in red. *D*, Methylation of the ZHX2 N-terminal peptide by METTL11A is significantly lower when METTL13 is present compared to when METTL11A is alone. METTL13 does not methylate the ZHX2 peptide. *E*, Methylation of the ZHX2 N-terminal peptide is significantly higher when METTL11B is present compared to when METTL11A is alone. When both METTL13 and METTL11B are present with METTL11A, methylation activity is significantly reduced compared to when METTL11A is alone. *F*, There is no significant difference between methylation activity of METTL11A against the RPS25 N-terminal peptide when METTL11B is present compared to METTL11A alone. Methylation activity is significantly lower when METTL13 is present with METTL11A, with or without METTL11B also present. These data indicate METTL11A, METTL11B, and METTL13 can exist in a complex together, and the regulatory effects of METTL13 on the methylation activity of METTL11A take precedence over the effects of METTL11B. Unless otherwise specified, n=3, * denotes p<0.05, ** denotes p<0.01, *** denotes p<0.001, **** denotes p<0.0001 as determined by unpaired t-test.

Finally, as it appears all three family members can exist in a complex together, we were interested in determining if the regulatory effects of METTL13 or METTL11B took precedence when all three enzymes were present. As METTL11B has previously been shown to activate the activity of METTL11A specifically for non-canonical substrates (4,12), we used *in vitro* methyltransferase assays to measure if METTL13 also affected the ability of METTL11A to methylate the non-canonical substrate ZXH2. We found that, similar to the canonical RCC1 substrate (Fig. 3A), METTL13 inhibited METTL11A trimethylation of a ZHX2 N-terminal peptide (Fig. 7D). We replicated previous findings that METTL11B significantly increased METTL11A methylation activity on the ZHX2 N-terminal peptide (4), but interestingly, found that inhibition by METTL13 takes precedence over activation by METTL11B. We found a significant decrease in ZHX2 methylation when both METTL11B and METTL13 are present compared to METTL11A alone (Fig. 7E). As ZHX2 is a nuclear METTL11A substrate, we also wanted to determine if METTL13 could inhibit METTL11A methylation of cytoplasmic targets in the presence or absence of METTL11B. Using the N-terminal peptide of the cytoplasmic, canonical substrate RPS25, we found METTL13 could inhibit METTL11A methylation of RPS25 in both the presence and absence of METTL11B (Fig. 7F). These results suggest that METTL13 inhibits methylation of both nuclear and cytoplasmic METTL11A substrates, and that these inhibitory effects are not altered by the presence of METTL11B. Together, these findings suggest a model where METTL13 is able to inhibit METTL11A in the cytoplasm, even if it translocates to the cytoplasm with METTL11B.

## Discussion

We had previously shown that METTL11B binds METTL11A and activates its methylation activity towards non-canonical substrates (12). Here we further refine our predicted model of the METTL11A/METTL11B complex and identify mutations that disrupt its interaction. We also verify that METTL11A participates in an additional regulatory interaction with METTL13 in the cytoplasm. For both the canonical and non-canonical substrates tested, METTL13 inhibits METTL11A activity, while METTL11A inhibits METTL13 methylation of the eEF1A N-terminus and promotes K55 methylation. These regulatory functions are distinct from their catalytic functions, as catalytically inactive mutants elicit levels of inhibition comparable to WT. Finally, we found that all three enzymes can exist in a complex together, with METTL11A likely acting as a bridge. When METTL11A, METTL11B, and METTL13 are together, METTL13 outcompetes METTL11B to inhibit METTL11A methylation activity.

These are the first studies to show a methyltransferase participating in opposing regulatory interactions with two different family members and identify novel, non-catalytic regulatory functions for both METTL11A and METTL13.

Together, these data suggest a model where cellular localization plays an important role in determining the interaction partners and main function of METTL11A. In the nucleus, METTL11A catalytic activity predominates and can be activated by METTL11B. In the cytoplasm, METTL11A is inactive against nuclear and cytoplasmic targets (3), due to inhibition by METTL13. Here, its non-catalytic role of METTL13 regulation predominates. It remains unknown if METTL11B can translocate out of the nucleus with METTL11A, but our data show that if this were the case, it would not affect the inhibitory effects of METTL13 on METTL11A. While we see very little METTL13 in nuclear fractions (Fig. 2B), there are reports of nuclear METTL13 (43). Our data suggest that if it enters the nucleus it will be able to inhibit METTL11A activity, so it will be interesting to determine under what conditions METTL13 translocates to the nucleus and if it is in concentrations high enough to inhibit METTL11A activity.

Methyltransferases taking on different roles in different cellular compartments has become an increasingly apparent trend. METTL3, the active subunit of the primary complex responsible for the N^6^-methyladenosine (m^6^A) modification, typically localizes to the nucleus, where it interacts with METTL14 and additional proteins to form the m^6^A writer complex (49,50). However, METTL3 has also been found in the cytoplasm where it interacts with different binding partners (translation initiation machinery) to enhance mRNA translation (51,52). Interestingly, METTL14, the principal activator of METTL3 activity, is only found in the nucleus (51). Another member of the METTL family, METTL16 has also recently been found to take on two distinct roles depending on cellular localization (44). Similar to METTL3, METTL16 in the nucleus places the m^6^A modification on a set of distinct RNA targets (53) and in the cytoplasm promotes translation through direct interactions with translation initiation machinery (44). Relative to METTL3, which is predominantly nuclear, METTL16 has a much larger cytoplasmic population, suggesting its role in the cytoplasm may be more impactful (44). It will be interesting to determine if this cytoplasmic role of METTL11A is also related to promoting translation through its ability to increase methylation of K55 eEF1A by METTL13.

Similar to METTL11A, the distinct nuclear versus cytoplasmic functions found in other METTL proteins can also be divided into catalytic versus non-catalytic functions, respectively. Both METTL3 and METTL16 were reported to maintain their cytoplasmic translation regulatory functions independently of their own catalytic activity (44,51). Again, our findings with METTL11A and METTL13 are consistent with this pattern in that their regulatory effects were also still elicited by catalytically inactive mutants. This pattern highlights a distinct division among the METTL family of proteins, as some methyltransferases have both catalytic and non-catalytic functions like METTL3, METTL16, METTL11A and METTL13, while others, like METTL14 and perhaps METTL11B, only seem to have non-catalytic functions. We suspect additional functions will be identified for methyltransferases that were previously thought to only have one defined role.

Since the regulatory interactions are independent of catalytic activity, it is likely that the described regulatory effects are the result of structural changes elicited through binding. Future studies will use nuclear magnetic resonance (NMR) to delineate structural changes occurring in METTL11A following the binding of either METTL11B or METTL13. By comparing the structural changes that occur in METTL11A when it interacts with its binding partners to the changes that occur with substrate binding, we predict that the binding of METTL11B may promote a METTL11A conformation that is more favorable for non-canonical substrate binding, subsequently activating methylation activity. An interaction with METTL13 on the other hand, may promote a conformational change in METTL11A that makes substrate binding unfavorable, thereby inhibiting its methylation activity.

The identification of these novel, non-catalytic functions of both METTL11A and METTL13 is important because despite the many similarities between the various members of the METTL family, each displays their own unique and complex characteristics surrounding their regulatory mechanism, localization pattern, and functions outside of their methylation activities. By having a more thorough understanding of each METTL protein, we can also learn about how their dysregulation, mislocalization, or misfunction can lead to the development of diseases. Disruptions to non-catalytic, regulatory functions are known to manifest as abnormal catalytic functions, as a mutation in METTL14 (R298P) leads to decreased methylation activity of the METTL3/METTL14 complex and also disrupts the ability of the complex to distinguish between the WT and mutant RNA substrate (14). R298 of METTL14 is located close to the METTL3/METTL14 junction and catalytic site, and this mutation is frequently found in human endometrial cancer, suggesting the biological relevance of a mutation in the inactive activator of the METTL3/METTL14 complex (14-16,35,54).

Future studies will look to identify important residues mediating the METTL11A/METTL13 interaction, as we have done with METTL11A/METTL11B. We suspect some of these interacting residues will also be found in human diseases, given the large number of mutations that have been found present in various cancer types for METTL11A, METTL11B, and METTL13 (35), which could possibly be eliciting problematic effects through the disruption of their non-catalytic functions. We suspect that as the interaction sites between METTL11A, METTL11B, and METTL13 become more defined, we will pinpoint additional biologically relevant mutations disrupting regulatory interactions and ultimately affecting Nα-methylation patterns. As rational design of peptidomimetic molecules that disrupt protein-protein interactions is a growing trend in drug development (55), targeting these methyltransferase interactions may also be a promising therapeutic option moving forward.

## Experimental Procedures

### Molecular cloning and recombinant protein purification

The full-length human METTL11A-FLAG, METTL11B-FLAG, and METTL11B-GFP constructs used in the co-immunoprecipitations (co-IPs) were cloned as previously described (1,2), and they were also used as templates for constructing the subsequent METTL11A-FLAG and METTL11B-GFP mutants using the Quikchange site-directed mutagenesis protocol (Agilent Technologies, Santa Clara, CA). The following forward primers and their reverse complements were used: Q67H METTL11B: 5’-GTCATCAATGGTGAGATGCATTTCTATGCCAGAGCTAAAC-3’; F68L METTL11B: 5’-GTCATCAATGGTGAGATGCAGTTGTATGCCAGAGCTAAAC-3’; V224L METTL11B: 5’-CATATTGAAGGACAATCTGGCCCGGGAGGGCTGTATC-3’; D232N METTL11B: 5’-GGGAGGGCTGTATCCTTAATCTCTCTGACAGCAGTGTGAC-3’; R78G METTL11A: 5’-GAGGATCACCAAGGGGCTGCTCCTGCC-3’. Human METTL13 (Horizon Discovery, Waterbeach, UK) was cloned into the XbaI and BamHI sites of pKGFP2 to remove one copy of GFP and create the METTL13-GFP construct for co-IPs. The truncated METTL13 constructs were subcloned from this full-length METTL13-GFP construct. METTL13 was also amplified to introduce a 5’XbaI restriction site, C-terminal Flag tag, and a 3’BamHI restriction site, and subcloned into pKH3 (a generous gift from Dr. Ian Macara, Vanderbilt University) to create the METTL13-FLAG construct used for mass spectrometry. The following primers were used: 5’XbaIMETTL13: 5’-GCTCTAGAATGAACCTCTTACCTAAAAG-3’; 3’BamHIMETTL13: 5’-GCGGATCCCACAATTTTCACCGTCTTGAG-3’; 3’BamHIFLAGMETTL13: 5’-GCGGATCCTCATTTATCATCATCATCTTTATAATCCACAATTTTCACCGTCTTGAG-3’; METTL13_1-344_ Reverse: 5’-GCGGATCCATACTGCTGACCTCGGTGAAG-3’; METTL13_345-699_ Forward: 5’-GCTCTAGAATGGAAAGCATGGACCACATCCA-3’.

For recombinant protein purification, full-length, human METTL11A was cloned into pET15b (Millipore Sigma, Burlington, MA) as described previously (2). Catalytically inactive mutant (D180K) METTL11A-pET15b was cloned as previously described (1). Full-length human RCC1 was cloned into pET30a (Millipore Sigma) as described previously (56). METTL13 was cloned into the NdeI and XhoI restriction sites of pET15b (Millipore Sigma), and human eEF1A1 (Horizon Discovery) was cloned into the NdeI and XhoI restriction sites of pET30a (Millipore Sigma). Quikchange mutagenesis (Agilent Technologies) was used to obtain the catalytically inactive double mutant, METTL13 G58R E524A-pET15b. All recombinant His-tagged proteins were purified as described previously (57). The primers used to create the METTL13-pET15b and eEF1A-pET30a constructs, and the forward primers plus their reverse complements used to create the METTL13 G58R E524A-pET15b construct are as follows: 5’NdeIMETTL13: 5’-GCCATATGAACCTCTTACCTAAAAGTTC-3’; 3’XhoIMETTL13: 5’-GCCTCGAGTCACACAATTTTCACCGTCT-3’; 5’NdeIeEF1A: 5’-GCCATATGGGAAAGGAAAAGACTCATATCAACATTGTCG-3’; 3’XhoIeEF1A: 5’-CCCTCGAGTTTAGCCTTCTGAGCTTTCTGGGC-3’; METTL13G58RQCF: 5’-TCTGAGTTGCGACACCCAATCACCAGCACC-3’; METTL13E524AQCF: 5’-CATGGAGGGATCGATCGCCACAGCATCAATGCA-3’

### Cell culture

Human embryonic kidney (HEK293T) cells were maintained in Dulbecco’s Modified Eagle Medium (DMEM, Corning Incorporated, Corning, NY) supplemented in 10% fetal bovine serum (FBS, R&D Systems, Inc., Minneapolis, MN) and 1% penicillin-streptomycin (P/S, Corning Incorporated). HCT116 human colorectal carcinoma and CRISPR/Cas9-mediated METTL11A knockout (KO) HCT116 cell lines were cultured in McCoy’s 5A Modified Medium (Lonza, Walkersfield, MD) supplemented with 10% FBS and 1% P/S. METTL11A KO cells were generated previously (10). Maintenance of METTL11A loss was verified by western blot (Fig. 7A). HEK293T and HCT116 cell lines were a generous gift from Dr. Ian Macara.

### Western blots

Protein concentrations were measured using the Pierce 660 nM Protein Assay (Thermo Fisher Scientific, Waltham, MA), and samples were normalized for equal protein loading prior to separation on 10% SDS-PAGE gels. Gels were transferred to nitrocellulose membranes using a Trans-blot Turbo Transfer System (Bio-Rad Laboratories, Inc., Hercules, CA). Membranes were blocked for 1 hour in 5% w/v nonfat dry milk in TBST (TBS + 0.1% Tween). Primary and secondary antibodies were also diluted in the 5% TBST-milk solution. Primary antibodies were used at the following dilutions: rabbit anti-GFP 1:1000 (Cell Signaling Technologies, Danvers, MA); rabbit anti-FLAG-HRP 1:500 (Millipore Sigma); goat anti-RCC1 1:1000 (Santa Cruz Biotechnology, Inc., Dallas, TX); rabbit anti-β-tubulin 1:1000 (Cell Signaling Technologies); rabbit anti-METTL11A 1:3000 (1); rabbit anti-eEF1A 1:1000 (Cell Signaling Technologies); and rabbit anti-dimethyl-eEF1A-K55 1:1000 (Abclonal, Woburn, MA). Secondary antibodies used were donkey anti-rabbit and donkey anti-goat at 1:5000 dilutions (Jackson ImmunoResearch, West Grove, PA). Blots were developed using either Clarity Western ECL Substrate (Bio-Rad Laboratories, Inc.) or SuperSignal West Femto Maximum Sensitivity Substrate (Thermo Fisher Scientific) and imaged on a ChemiDoc Touch imaging system (Bio-Rad Laboratories, Inc.). Blot quantifications were performed using ImageJ 1.52a (NIH, Bethesda, MD).

### Co-immunoprecipitation (co-IP) experiments

Twenty-four hours prior to transfection, 1×10^6^ HEK293T cells or 4×10^6^ HCT116 control or METTL11A KO cells were plated in 10 cm tissue culture dishes. HEK293T cells were calcium phosphate-transfected with 1 µg each of appropriate constructs, and HCT116 cells were transfected with 8 µg each of appropriate constructs using Lipofectamine 2000 (Thermo Fisher Scientific). Approximately 24 hours post-transfection, cells were scraped directly into 200 µL of lysis buffer (50 mM Tris, 300 mM NaCl, 5 mM MgCl_2_, 1% NP-40, 7 mM BME), plus protease inhibitors. Twenty µL of cell lysate was saved for input controls. The remainder of the lysate was added to 5 µL of Pierce Protein G agarose beads (Thermo Fisher Scientific), and the mixture was rotated 1-2 hours at 4°C to pre-clear. Following the pre-clear incubation, the mixtures were spun quickly, and the super was added to 40 µL of EZ View Red anti-FLAG M2 agarose beads (Millipore Sigma). The mixture was rotated 1-2 hours at 4°C and washed 3x with PBS + 0.1% NP-40 + 500 mM NaCl. The immunoprecipitated proteins were eluted from the beads in 5x Laemmli buffer and boiled at 95°C for 10 mins. The bead-free IP supernatant and the input samples were run on 10% SDS-PAGE gels and analyzed with western blots as described above.

### *In vitro* methyltransferase assays

*In vitro* methyltransferase assays for measuring Nα-methylation were conducted using the MTase-Glo Methyltransferase Assay (Promega, Madison, WI) following the manufacturer’s protocol. Each assay used 0.2 µM recombinant enzyme unless otherwise specified for the titration of METTL11A, 20 µM S-adenosyl methionine (SAM), and either 1 µM of RCC1 (SPKRIAKRRSPPADA), 10 µM of eEF1A (GKEKTHINIVVIGH), 60 µM of ZHX2 (ASKRKSTTPCMVRTS) or 0.75 µM of RPS25 (PPKDDKKKKDAGKS) N-terminal peptides (Anaspec, Fremont, CA) as substrates. Briefly, reactions between the various combinations of enzymes and substrates were incubated in wells on a 96-well plate at room temperature for either 5 or 20 minutes and stopped with the addition of 0.5% trifluoroacetic acid. The MTase-Glo detection reagents were added according to the manufacturer’s protocol, and the luminescence was measured using the Cytation5 Imaging System (BioTek, Winooski, VT). Background signals were measured through the inclusion of no substrate control reactions and subtracted where applicable.

The *in vitro* methyltransferase assays to measure full-length, recombinant eEF1A1 K55 methylation and RCC1 Nα-methylation were conducted using reactions consisting of 2 µg recombinant METTL13 and/or 2 µg recombinant METTL11A, 3 µg recombinant eEF1A1 or 2 µg recombinant RCC1 substrate, and 100 µM SAM. The reaction volume was adjusted to 50 µL with methyltransferase buffer (50 mM potassium acetate, 50 mM Tris/HCl, pH 8.0). The reactions were incubated at 30°C for 10 minutes, with 15 µL aliquots taken at 0, 5 and 10 minutes. Reaction aliquots were mixed with 5x Laemmli buffer and analyzed by western blotting.

### Nuclear/cytoplasmic fractionations and mass spectrometry

HEK293T cells were transfected as described previously to express METTL13-FLAG. Twenty-four hours post-transfection, lysates from METTL13-FLAG expressing cells and untransfected control cells were fractionated to yield both a nuclear and a cytoplasmic fraction. This process was repeated for four 10 cm plates for each condition, either METTL13-FLAG or untransfected. Briefly, 24 hours post-transfection, cells were collected directly into cytoplasmic lysis buffer (10 mM HEPES, 1.5 mM MgCl_2_, 10 mM KCl, 0.5 mM DTT, 0.05% NP-40, pH 7.9) plus protease inhibitors. To disrupt the cellular membrane, lysates were vortexed on high ∼7 seconds, incubated on ice 10 minutes, vortexed again, incubated on ice 1 minute, then spun down at high speed (13,200 rpm) for 5 minutes at 4°C. Supernatant was collected as the cytoplasmic fraction. Remaining nuclei were washed 3x with cold PBS and resuspended in a nuclear lysis buffer (5 mM HEPES, 1.5 mM MgCl_2_, 0.2 M EDTA, 0.5 mM DTT, 26% glycerol (v/v), 300 mM NaCl, pH 7.9). Nuclei were ruptured by vortexing on high for 15 seconds, and incubating on ice for 40 minutes, with a 15 second vortex every 10 minutes. Supernatant was collected following a 10-minute spin at 4°C (13,200 rpm) as the nuclear fraction. Fractionation cleanliness was tested by analyzing input samples (20 µL of lysate) by western blot with RCC1 (nuclear protein) as a readout of cytoplasmic fraction cleanliness, and tubulin (cytoplasmic protein) as a readout of nuclear fraction cleanliness (Fig. 2B).

The remainder of the cytoplasmic fraction was added to 5 µL of washed Pierce Protein G agarose beads (Thermo Fisher Scientific), and the mixture was rotated 1-2 hours at 4°C to pre-clear. Following the pre-clear, the mixture was spun quickly, and the super was added to 15 µL of washed Pierce anti-DYKDDDDK (FLAG) magnetic agarose beads (Thermo Fisher Scientific). The mixture was rotated 1-2 hours at 4°C and washed 3x with PBS + 0.1% NP-40 + 150 mM NaCl. Interacting proteins were eluted into buffer (25 mM HEPES-NaOH, pH 7.5, 100 mM NaCl, 0.1 mg/mL FLAG peptide (Anaspec), and 10 µL/mL PMSF, filter sterilized). Elution was sent for label-free quantification mass spectrometry analysis at the Cornell Institute of Biotechnology. Top interactors were those 100% abundant in METTL13-FLAG sample compared to untransfected control.

### Molecular modeling

Molecular modeling was produced using a combination of the ZDOCK server, the I-TASSER server, and Chimera UCSF (36,37,46-48). The METTL11A dimer model was produced using the METTL11A crystal structure of the monomer (PDB: 2EX4, chain A) as both input structures in the protein-protein docking tool, ZDOCK (36). Out of the top 10 predicted models, the one most closely resembling the previously published METTL11A dimer model (12) was selected for use in further modeling. To produce the METTL11A/METTL11B heterotrimer model with the METTL11B D232 constraint, the METTL11A dimer and the METTL11B crystal structure (PDB: 5DUB) were used as inputs in ZDOCK (36), with METTL11B D232 selected as a contacting residue. To obtain a predicted structure of the METTL13 N-terminal domain (M1-Y344), the I-TASSER server was used (46-48). This predicted structure of the METTL13 N-terminal domain was then used as input along with the METTL11A dimer model in ZDOCK to obtain a model to predict the METTL11A/METTL13 interaction interface (36,46-48). Interacting residue pairs predicted to be involved in the interaction were selected based on having a distance between interacting residues of less than 4 Å, satisfying the maximum Van der Waals radius.

Molecular graphics and analyses were performed with UCSF Chimera developed by the Resource for Biocomputing, Visualization, and Informatics at the University of California, San Francisco, with support from NIH P41-GM103311 (37).

### Statistical analyses

All statistical analyses were performed using GraphPad Prism 6 software (GraphPad, San Diego, CA). The specific statistical tests used are noted in the respective figure captions, and results are presented as mean ± standard error of the mean (SEM).

## Data availability

All data is contained within the manuscript.

### Supporting information

This article contains supporting information.

## Acknowledgments

We thank Schaner Tooley lab members James Catlin, Meghan Conner, Joseph Dombrowski, and John Tooley for their critical reading of the manuscript. Mass spectrometry was performed by the Proteomics and Metabolomics Facility (RRID:SCR 021727) at the Cornell Institute of Biotechnology.

## Author contributions

H.P. performed all the experiments and data analysis, wrote the original draft, and prepared the figures. C.S.T conceptualized and designed the project and provided supervision and editing.

## Supporting Information

**Supplemental Figure 1.**
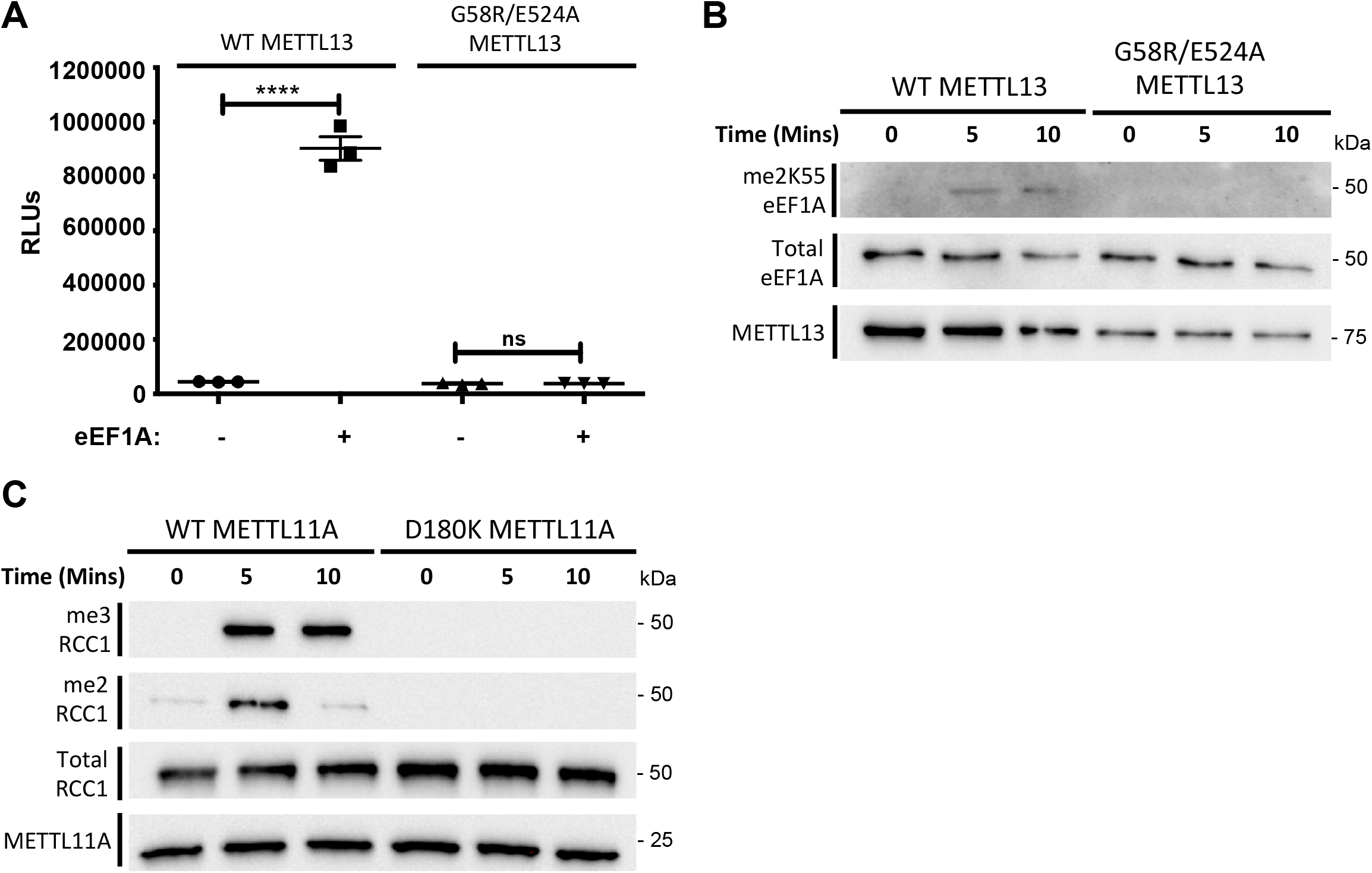
G58R/E524A METTL13 and D180K METTL11A are catalytically inactive. *A*, G58R/E524A METTL13 is unable to significantly methylate the N-terminal eEF1A peptide. *B*, G58R/E524A METTL13 is unable to methylate K55 eEF1A. *C*, D180K METTL11A is unable to methylate RCC1. **** denotes p<0.0001 as determined by unpaired t-test.

